# A novel mechanism of ubiquitin-charging a bivalent E2:RING E3 complex

**DOI:** 10.64898/2026.09.04.749512

**Authors:** Samuel Perreault, Ayush Mistry, Edwin Antony, Mark Hedglin

## Abstract

Protein ubiquitination results from a cascade of enzyme interactions that transfer ubiquitin from one covalent bond to another. First, an E1 is activated by attaching ubiquitin to itself through a thioester bond. Next, in E2 charging, activated E1 interacts with an E2 and transfers ubiquitin to a thioester bond on the E2. Finally, an E3 mediates the transfer of ubiquitin from a charged E2 to an isopeptide bond on a protein target. RING E3s, the largest E3 family in eukaryotes, function as scaffolds rather than enzymes and facilitate direct transfer of ubiquitin from a charged E2 to a protein target by simultaneously engaging both. All RING E3s engage an E2 at an interface that overlaps with the E1-binding site and several form stable, bivalent complexes with an E2 through additional binding surfaces. How E2 charging proceeds within these complexes remains unclear. Here, we utilize human Rad6(Rad18)_2_ as a model bivalent E2:RING E3 complex to delineate the interplay of protein•protein interactions among the primary human E1 (Uba1), an E2 (Rad6), and a RING E3 (Rad18) during E2 charging. Collectively, the results reveal a novel mechanism that is not directed by ubiquitin thioester “affinity switches.” Rather, interactions of Rad18 with Rad6 slow the chemistry step and all preceding steps of the Uba1 catalytic cycle via competitive inhibition but accelerate release of charged Rad6 from Uba1, ultimately stimulating Rad6 charging overall. To the best of our knowledge, this represents the first example of a RING E3 stimulating Uba1-dependent charging of an E2.

## Introduction

Protein ubiquitination is a post-translational modification (PTM) involved in nearly all processes of human biology and is the result of a cascade of enzyme interactions that transfer ubiquitin from one covalent bond to another. The enzymes involved are referred to as ubiquitin-activating (Uba/E1s), ubiquitin-conjugating (Ubc/E2s), and ubiquitin ligases (E3s). The cascade begins with the generation of an intermediate (Ub∼E1) in which ubiquitin is covalently attached to the active site cysteine of the E1 by a high-energy thioester bond. This step is ATP-dependent and referred to as E1 activation. Next, Ub∼E1 engages an E2 and transfers the thioester-linked ubiquitin to the active-site cysteine of the E2 via isoenergetic transthioesterification, forming Ub∼E2. This step is referred to as E2 charging. E3s mediate the final transfer of Ub from a charged E2 to the substrate through mechanisms that differ between E3 families. RING E3s represent the largest family of ubiquitin ligases in all eukaryotes and do not form Ub∼E3 thioester intermediates during protein ubiquitination. Rather, RING E3s function as scaffolds that simultaneously engage a charged E2 and a protein substrate, facilitating the direct transfer of ubiquitin from a high-energy thioester bond on the Ub∼E2 to a stable isopeptide bond on the protein substrate^1,2^.

Central to all ubiquitination cascades is the timely formation of charged E2:E3 complexes, which requires the intimate coordination of protein•protein interactions among E1, E2, and E3 binding partners. Uba1, the primary E1 in humans, engages a highly conserved E2 interface consisting of a residue within Loop 1, a residue within Loop 2, and three basic residues within Helix 1 that correspond to (K/R)RXX(K/R) and assume a triskelion of positive charge at one end of the overall E2 structure^3^. Interestingly, all RING E3s utilize a RING domain to engage an E2 interface that significantly overlaps with the aforementioned Uba1-binding site. Hence, E2 interactions with Uba1 and a RING domain are mutually exclusive, necessitating that an E2 disengage from the RING interface for ubiquitin transthioesterification by Uba1^1,3-7^.

Furthermore, several RING E3s, such as Rad18, brandish an additional E2-binding domain that engages an interface distinct from the Uba1-binding site, resulting in tight E2:RING E3 complexes stabilized by multiple E2-binding interactions. Here, we refer to these E2:RING E3 complexes as bivalent.

The RING E3 Rad18 functions only as a homodimer and is responsible for the attachment of single ubiquitin moieties (i.e., monoubiquitination) to PCNA, an essential DNA replication factor, and this PTM is imperative for cell recovery from DNA replication stress^8^. In addition to an N-terminal RING domain that engages the Uba1-binding site of its cognate E2, Rad6, Rad18 also contains a C-terminal Rad6-binding (R6B) domain that engages the noncovalent ubiquitin interaction site on the “backside” of Rad6, opposite the active-site cysteine residue (C88). This interaction prevents ubiquitin chain formation by Rad6 and is required for PCNA monoubiquitination^1,4-6,9-11^. Rad18 RING and R6B domains engage Rad6 independently of one another and together form a stable, bivalent interaction with Rad6^6,12-15^. However, Rad18 dimerization is asymmetric such that only one R6B domain and one RING domain are available for interaction with Rad6. Hence, the Rad18 homodimer, i.e., (Rad18)_2_, engages a single Rad6^6,11^.

Currently, it is unclear how charging occurs for bivalent E2:RING E3 complexes, limiting our fundamental understanding of RING E3-dependent ubiquitination cascades. In particular, it is unknown how the protein•protein interactions among an E1, an E2, and a RING E3 are coordinated in this process. In the present study, we address this knowledge gap by characterizing: 1) the stability of the native Rad6(Rad18)_2_ complex; 2) the kinetics of Rad6 charging by Uba1 in the absence and presence of the native, full-length Rad18 homodimer; and 3) the affinities of full-length (Rad18)_2_ for charged and uncharged (i.e., discharged) Rad6. Collectively, the results from these comprehensive studies reveal an overall novel mechanism of charging a bivalent E2:RING E3 complex that is not directed by ubiquitin thioester “affinity switches.” Rather, (Rad18)_2_ stimulates Rad6 charging by accelerating release of the Ub∼Rad6 product from Uba1.

## Experimental Procedures Reagents and Materials

Creatine phosphokinase from rabbit muscle Type I, salt-free, lyophilized powder, phosphocreatine disodium hydrate, bovine thrombin, and pepstatin A were purchased from MilliporeSigma (Sigma-Aldrich Inc., Saint Louis, MO). Aprotinin and leupeptin were purchased from VWR (Radnor, PA).

### Recombinant Human Proteins

Ubiquitin labeled with a single Cyanine5 (Cy5) moiety at a specific site (Cy5-Ub) was purchased from South Bay Bio (San Jose, CA). Cy5-Ub has a fully functional C-terminus and all lysine residues are available, allowing for further conjugation into ubiquitin chains. The Cy5-labeled ubiquitin does not affect E1-E2-E3 ubiquitin transfer/conjugation cascades (per South Bay Bio, San Jose, CA). Cy5-Ub is >98% pure (by LC-MS). The concentration of Cy5-Ub was verified using the extinction coefficient for Cy5 (ε_650_ = 250,000 M^-1^cm^-1^). N-terminal 6His-tagged human ubiquitin (His-Ub) was purchased from MilliporeSigma (Sigma-Aldrich Inc, Saint Louis, MO). N-terminal 6His-tagged human Uba1 was obtained via in-house preparations (described below) or purchased from either R&D Systems (Minneapolis, MN) or South Bay Bio (San Jose, CA). Label-free (i.e., native) human Uba1 was purchased from either R&D Systems (Minneapolis, MN) or South Bay Bio (San Jose, CA). SDS-PAGE analyses of in-house preparations of recombinant human proteins used in the present study are provided in **Figure S1**.

In-house preparations of N-terminal 6His-tagged human Uba1 were obtained by slight modifications to published protocols^13,16-18^. Specifically, immobilized metal affinity chromatography was updated from batch binding to cobalt resin to automated FPLC purification using HiTrap Talon columns (5 × 5 mL). Wash conditions were adjusted to incorporate defined steps ranging from 150–500 mM NaCl. Bound protein was eluted with a linear gradient of imidazole (20–500 mM) in buffer containing 150 mM NaCl. All subsequent purification steps were performed exactly as described previously^13,17,18^ with the exception that glycerol was included in the respective buffers to improve protein stability and purity. The absolute concentration of each frozen in-house preparation was determined by Bradford analysis of Coomassie-stained polyacrylamide gels using BSA as the standard. The active concentrations of all Uba1 samples were determined via modifications to published protocols, as described in the **Supplemental Information**^3,19-22^. For all Uba1 samples used in the present study, the concentration of active Uba1 was typically ∼50% of the absolute concentration of Uba1 (i.e., Uba1 was ∼50% active, **Figure S2**).

N-terminal 6His-tagged human Rad6 was purified via slight modifications to published protocols^13,18^. Specifically, induction was performed using an increased IPTG concentration of 0.5 mM, gravity-flow Ni-NTA resin was replaced with automated FPLC purification using a HisTrap FF column, and bound protein was eluted using 300 mM imidazole. All subsequent purification steps were performed as described previously^13,18^. The N-terminal 6His tag on recombinant human Rad6 was removed by incubation with bovine thrombin at a ratio of 5 units per mg Rad6 for 2 h at room temperature on a tube rotator. The cleavage reaction was loaded at 1 mL/min onto tandem 1 mL HiTrap Heparin HP and 1 mL HisTrap HP columns that were connected in succession and equilibrated in Rad6 storage buffer (50 mM HEPES•KOH, pH 7.5, 150 mM NaCl, 0.5 mM TCEP, 10% glycerol v/v). Native Rad6 (i.e., tag free) was collected in the initial 10 mL of the flow-thru and concentrated via 3 kDa molecular weight cutoff filter device centrifugation. The concentration of native Rad6 was first determined from the calculated extinction coefficient (from ExPASy ProtParam) and subsequently verified via Coomassie-stained gel Bradford assay using BSA as a standard.

The pET28a plasmid encoding N-terminal 6His-SUMO-tagged human Rad18^23^, referred to herein as SUMO-Rad18, was transformed into *E. coli* Rosetta 2 (DE3) cells by heat shock and selected on LB agar containing kanamycin and chloramphenicol. Overnight starter cultures grown in LB with antibiotics were used to inoculate 12 L of LB medium (1:100 dilution).

Cultures were grown at 37 °C to an OD600 of 0.5–0.6, shifted to 15 °C, and induced with 0.5 mM IPTG in the presence of 100 µM ZnCl_2_. Cultures were grown for an additional 8 h at 15 °C. Cells were harvested by centrifugation at 11,000 × g for 20 min at 4 °C, and the pellet was not frozen prior to lysis. Cell pellets were resuspended at approximately 4 mL per gram of pellet in lysis buffer (10 mM HEPES•KOH, pH 7.5, 0.5 mM TCEP) supplemented with 150 mM NaCl, leupeptin (1 µg/mL), aprotinin (1 µg/mL), pepstatin A (1 µg/mL), and PMSF (1 mM). Cells were lysed by sonication while maintaining the lysate temperature below 9 °C, and the resultant lysates were clarified by two sequential centrifugation steps at 26,000 × g for 20 min at 4 °C. The clarified lysate was applied to a 15 mL (3 × 5 mL) HiTrap Talon column equilibrated in lysis buffer supplemented with 150 mM NaCl. The column was washed sequentially with 10 column volumes (CV) each of lysis buffer supplemented with the following: 5 mM imidazole and 150 mM NaCl; 5 mM imidazole and 500 mM NaCl; 10 mM imidazole and 300 mM NaCl; and 20 mM imidazole and 150 mM NaCl. Bound protein was eluted with a 10 CV linear gradient of imidazole (20–500 mM) in lysis buffer supplemented with 150 mM NaCl. Fractions containing full-length protein were pooled and applied directly to a 5 mL HiTrap Heparin column equilibrated in lysis buffer supplemented with 150 mM NaCl. The column was washed sequentially with 10 CV each of lysis buffer supplemented with the following concentrations of NaCl: 150 mM; 200 mM; 300 mM; and 400 mM. Bound protein was eluted with a 10 CV linear gradient of NaCl (400 mM to 1 M) in lysis buffer. Fractions containing full-length protein were pooled, concentrated by centrifugation in a 30 kDa molecular weight cutoff centrifugal filter device, divided into aliquots, flash-frozen in liquid nitrogen, and stored at −80 °C. The concentration of the frozen protein stock was determined via Coomassie-stained gel Bradford assay using BSA as a standard. The His-SUMO tag on Rad18 is required to purify soluble protein in the absence of Rad6 and does not compromise Rad18’s functions with Rad6^6,9,18^.

Human Rad6(Rad18)_2_ containing Rad18 with an N-terminal 6His tag was obtained by modifications to published protocols, as follows^6^. Rad6 and Rad18 were co-expressed from plasmids (pCDF1b-Rad6b and pET28-Rad18) provided by Dr. Titia Sixma (Netherlands Cancer Institute, The Netherlands) in BL21(DE3) cells grown in LB medium supplemented with kanamycin and streptomycin sulfate. Cells were lysed in Tris•HCl buffer (pH 8.0) containing 300 mM NaCl, 2 µM ZnCl_2_, 5 mM imidazole, and 0.5 mM TCEP, in the presence of EDTA-free protease inhibitor tablets (1 tablet per 25 mL of buffer) and 1 mM PMSF. Immobilized metal affinity chromatography was transitioned from batch binding with Talon beads to automated FPLC purification using HiTrap Talon columns (3 × 5 mL), with defined binding and wash conditions ranging from 5–25 mM imidazole and 150–300 mM NaCl, followed by elution with 200 mM imidazole. Heparin chromatography was performed using a 50 mM NaCl wash followed by linear elution to 600 mM NaCl, and selective pooling of heparin-eluted peaks enabled removal of the majority of truncated Rad18 species present in the complex. Final purification was performed by size-exclusion chromatography on Superdex 200 using Tris•HCl buffer (pH 8.0) containing 150 mM NaCl and 0.5 mM TCEP, which allowed for further removal of truncated Rad18-containing species. The concentration of native Rad6 within Rad6(Rad18)_2_ was determined via Coomassie-stained gel Bradford assay using native Rad6 as a standard. The active concentration of Rad6, either as free Rad6 or Rad6(Rad18)_2_, was determined via modifications to published protocols, as described in the **Supplemental Information**. The concentration of active Rad6 used in the assays of the present study, either as free Rad6 or Rad6(Rad18)_2_, is equal to the absolute concentration of Rad6 (i.e., Rad6 is 100% active) (**Figure S3**).

Rad6^C88K^-Ub isopeptide conjugates were prepared by slight modifications to published protocols^24,25^. First, the active-site cysteine (C88) of human Rad6 was mutated to lysine via site-directed mutagenesis of the pET28A expression plasmid encoding N-terminal 6His-tagged human Rad6^13,18^ using the Q5® Site-Directed Mutagenesis Kit (New England Biolabs, Ipswich, MA) and the mutagenic primers below, which were designed with NEBaseChanger (New England Biolabs, Ipswich, MA).

Forward: 5’-TGGTAGCATAAAGTTAGATATCCTGCAG-3’ Reverse: 5’- TCAGCATACACATTTGGATG-3’

Following PCR amplification and transformation into chemically competent *E. coli*, individual colonies were selected, plasmid DNA was extracted and purified using the E.Z.N.A. Plasmid DNA Mini Kit (Omega Bio-Tek), and site-directed mutagenesis was verified by DNA sequencing via Plasmidsaurus Inc. (Louisville, KY). N-terminal 6His-tagged human Rad6^C88K^ was purified and the N-terminal 6His tag was removed, exactly as described above for Rad6. The concentration of the resultant protein (Rad6^C88K^) was first determined from the calculated extinction coefficient (from ExPASy ProtParam) and subsequently verified via Coomassie-stained gel Bradford assay using native Rad6 as a standard. Next, 60 µM Rad6^C88K^ was incubated with 120 µM 6His-Ub and 4 µM N-terminal 6His-tagged human Uba1 at 35 °C for 24 h in buffer containing 50 mM Tris•HCl, pH 10.0, 150 mM NaCl, 5 mM MgCl_2_, 0.8 mM TCEP, and 10 mM ATP. This reaction yields Rad6^C88K^-Ub-His conjugates in which the C-terminus of His-Ub is covalently linked to lysine at position 88 of human Rad6^C88K^ via an isopeptide bond.

Following completion of the reaction, Rad6^C88K^-Ub-His was first separated from unconjugated Rad6^C88K^ via HisTrap HP affinity column chromatography, and then Rad6^C88K^-Ub-His was isolated from His-Ub via size-exclusion chromatography. The concentration of Rad6^C88K^-Ub-His, referred to herein as Rad6^iso^-Ub, was first determined from the calculated extinction coefficient (from ExPASy ProtParam) and subsequently verified via Coomassie-stained gel Bradford assay using Rad6^C88K^ as a standard.

### Mass Photometry

All measurements were carried out on the TwoMP instrument (Refeyn Ltd., Oxford, United Kingdom). Glass coverslips (No. 1.5H thickness, 24 × 50 mm, VWR, Radnor, PA) were cleaned, dried, and prepared on the objective lens (Olympus PlanApo N, 1.42 NA, 60X), as described previously^26^. All dilutions and measurements were performed at room temperature (23 ± 2 °C) in 1X ubiquitin transfer buffer (25 mM HEPES•KOH, pH 7.5, 10 mM Mg(OAc)_2_, 100 mM KOAc) supplemented with 0.1 mM DTT. The final ionic strength was adjusted to physiological levels (200 mM) by adding appropriate amounts of KOAc. Rad6(Rad18)_2_ samples were allowed to equilibrate for 5 min, after which 1 µL of the respective sample was quickly diluted onto the sample stage, and video recording commenced and continued for 1 min. High-contrast (light-scattering) events corresponding to single-particle landings on the coverslip were identified and analyzed further. A known mass standard (β-amylase, SIGMA A8781-1VL) was used to convert the image contrast signal into mass units. Histograms were generated from all the data gathered during the 1 min video interval and fit to a single Gaussian function using non-linear least squares, as described previously^26^.

### Assays monitoring the formation of ubiquitin thioester conjugates

All assays were performed at room temperature (23 ± 2 °C) in 1X ubiquitin transfer buffer supplemented with 1X ATP regeneration system (10 mM phosphocreatine disodium hydrate, 10 units/mL creatine phosphokinase) and 0.1 mM DTT. The final ionic strength was adjusted to physiological levels (200 mM) by adding appropriate amounts of KOAc. All samples containing Cy5 were protected from light whenever possible. Reaction aliquots (2 µL) were removed at the indicated time points and quenched into 58 µL of 1.034X quench buffer, resulting in a 30-fold dilution of the reaction and a final 1X quench buffer condition (50 mM Tris•HCl, 10% glycerol, 2% SDS, 175 mM EDTA). After all time points were completed, samples were centrifuged at 21,100 × g for 5 min. A 2 µL aliquot of each quenched sample was resolved on a 15-lane, 4– 20% Mini-PROTEAN® TGX− Gel (Bio-Rad, Hercules, CA) run at 150 V. Under the gel running conditions, the tracking dye (bromophenol blue) typically used to track the progression of small proteins through the gel overlaps with Cy5-Ub and is partially observed when imaging. To remove interference from tracking dye in analyzed samples, the following precautions were employed. First, tracking dye was omitted from all quenched samples. Second, gels were loaded and run in a manner that tracked progression of the samples while also allowing the tracking dye to run into the lower buffer chambers. Specifically, all gels were first pre-run for 6 min at 150 V, tracking dye was added to the middle lane (lane 8) only, and the gel was run for an additional 6 min at 150 V. Finally, experimental samples (lacking tracking dye) were loaded in lanes 1–6 and 10–15, and the gels were run at 150 V until the tracking dye (in lane 8) ran off the bottom of the gel and into the lower buffer chamber. Resolved gels were then rinsed three times with ddH_2_O, imaged, and analyzed as described previously^13,17,18^. For visualization of the representative fluorescence scans presented in the **Supplemental Information** (see below), lanes 7–9 are removed via Photoshop editing of the original scans. For reference, representative fluorescence scans from single-turnover kinetic assays performed with Rad6 and Rad6(Rad18)_2_ (described below) are displayed in **Figure S4**.

The transfer of ubiquitin from Uba1 to Rad6 via transthioesterification (i.e., Rad6 ubiquitin thioester formation) was monitored in an assay adapted from previous procedures (**Figure 1**)^3,19-22,27^. Each of the kinetic assay conditions described below differs only by the respective concentrations of active Uba1 and active Rad6, with Rad6 present either as free Rad6 or Rad6(Rad18)_2._ For all assays, ATP (1 mM) was first pre-incubated with a 1X ATP regeneration system to generate a uniform ATP solution. Next, Uba1 and Cy5-Ub (5 µM) were added in succession, and the resultant solution was pre-incubated to form stoichiometric Cy5-Ub∼Uba1 thioester conjugates (where “∼” denotes a high-energy, thioester covalent bond) that are noncovalently bound to an acyl-adenylated ubiquitin. For simplicity, acyl-adenylated ubiquitins engaged by Cy5-Ub∼Uba1 thioester conjugates are omitted from all schematic representations.

**Figure 1.**
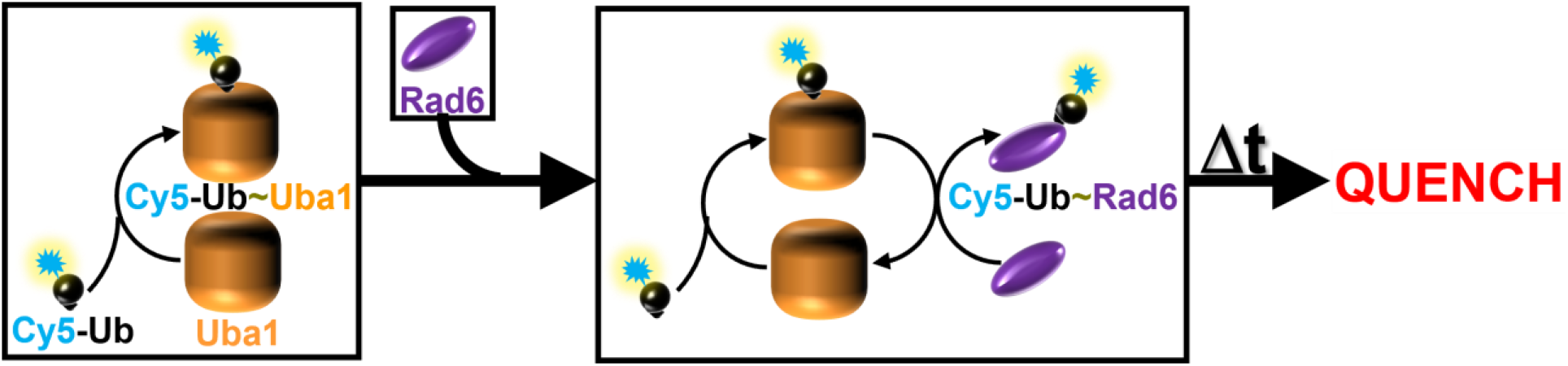
Schematic representation of the general kinetic assay to monitor Rad6 ubiquitin thioester formation. Preformed Cy5-Ub∼Uba1 thioester conjugates transfer Cy5-Ub to Rad6 residue C88 via a high-energy, thioester covalent bond, i.e., ubiquitin transthioesterification. “∼” denotes a high-energy, thioester covalent bond.

Herein, the formation of stoichiometric Cy5-Ub∼Uba1 thioester conjugates is referred to as activation. Also, Cy5-Ub∼Uba1 thioester conjugates and Uba1 are referred to as activated Uba1 and apo Uba1, respectively. Finally, the reaction is initiated by the addition of Rad6, either as free Rad6 or Rad6(Rad18)_2_, and the concentrations of Cy5-Ub, activated Uba1, and Rad6 ubiquitin thioesters are monitored simultaneously over time via fluorescence image scanning, as described above^13,17,18^. Herein, the formation of Cy5-Ub∼Rad6 conjugates is referred to as charging, and Cy5-Ub∼Rad6 thioester conjugates are referred to as charged Rad6. Single-turnover assays were performed with ≥ 500 nM active Uba1 and 250 nM active Rad6 or Rad6(Rad18)_2_. Pre-steady state (i.e., “burst”) assays were performed with 2.5 nM active Uba1 and 200–800 nM of either active Rad6 or Rad6(Rad18)_2_. Steady state assays were performed with 0.25 nM active Uba1 and 200 nM active Rad6 or Rad6(Rad18)_2_. All pre-steady state and steady state assays monitored only ≤ 10% and ≤ 5% of the reaction progress, respectively, based on substrate depletion, such that product release is irreversible^28^. For direct comparisons of Rad6 and Rad6(Rad18)_2_ in each kinetic assay condition, the respective experiments were carried out with the same preparations of ATP, 1X ATP regeneration system, Cy5-Ub, and Uba1.

### Isothermal Titration Calorimetry (ITC)

SUMO-Rad18, Rad6, and Rad6^iso^-Ub were prepared in buffer containing 50 mM Tris•HCl, pH 7.5, 150 mM NaCl, and 0.5 mM TCEP before analysis. ITC data were collected on a TA Instruments Low Volume Auto Affinity ITC at 20 °C with a stirring rate of 125 rpm. The reaction cell contained SUMO-Rad18 at approximately 9 µM protomer concentration, and the syringe contained either Rad6 at approximately 35 µM or Rad6^iso^-Ub at approximately 34 µM. Three independently paired experiments were performed, with Rad6 and Rad6^iso^-Ub titrations conducted using the same SUMO-Rad18 preparation on the same day within each replicate. For each experiment, 20 injections of 3.0 µL were performed with 180 s spacing between injections. The first injection was excluded from the fitting of the integrated binding isotherm. Control titrations of Rad6 or Rad6^iso^-Ub into buffer were performed to assess heats of dilution and were used for background correction of the corresponding SUMO-Rad18 binding experiments.

Binding isotherms were fit to an independent one-site binding model using NanoAnalyze software to determine the dissociation constant (K_D_) and stoichiometry (i.e., Rad6:Rad18 ratio).

## RESULTS

### The Rad18 homodimer engages Rad6 in a tight complex

Multiple qualitative studies from independent laboratories, including our own, have noted the innate stability of human Rad6(Rad18)_2_ complexes^6,11-15^. Based on the individual affinities of Rad6 for an isolated Rad18 RING domain (35 µM) and an isolated Rad18 R6B domain (62 µM), the overall dissociation constant, K_D_, of the human Rad6(Rad18)_2_ complex is predicted to be ∼22 nM according to an additive binding energy model without linker constraints (**see Supplemental Information**)^6,9,29^. To assess the validity of this model and confirm the composition of Rad6(Rad18)_2_ under the kinetic assay conditions described below, we analyzed dilutions of Rad6(Rad18)_2_ by mass photometry (**Figure 2**). At the highest concentration analyzed (**Figure 2A**), nearly all of the sample (94%) behaves as a complex with an estimated molecular weight distribution (136 ± 42 kDa) that agrees very well with the calculated molecular weight of the Rad6(Rad18)_2_ complex (134.13 kDa). As the sample is diluted, near-complete shifts to a lower molecular weight complex are observed. Specifically, in the penultimate dilution (**Figure 2C**), 87% of the sample has a molecular weight distribution of 112 ± 69 kDa. At the most dilute concentration (**Figure 2D**), 91% of the sample has a molecular weight distribution of 115 ± 47 kDa. These values agree very well with the calculated molecular weight of the Rad18 homodimer alone (116.24 kDa), indicating that Rad6 is predominantly disengaged from (Rad18)_2_ only when the Rad6(Rad18)_2_ complex is diluted to a very low concentration (≤ 12.5 nM). Collectively, these results indicate that human Rad6(Rad18)_2_ is a tight, stable complex with an affinity in the low nM range, in agreement with the aforementioned model, and remains completely intact at moderate concentrations (≥ 62.5 nM). In all kinetic assays described below for Rad6(Rad18)_2_, the concentration of the complex is ≥ 200 nM to ensure that all Rad6 is engaged by Rad18 homodimers prior to incubation with activated Uba1.

**Figure 2.**
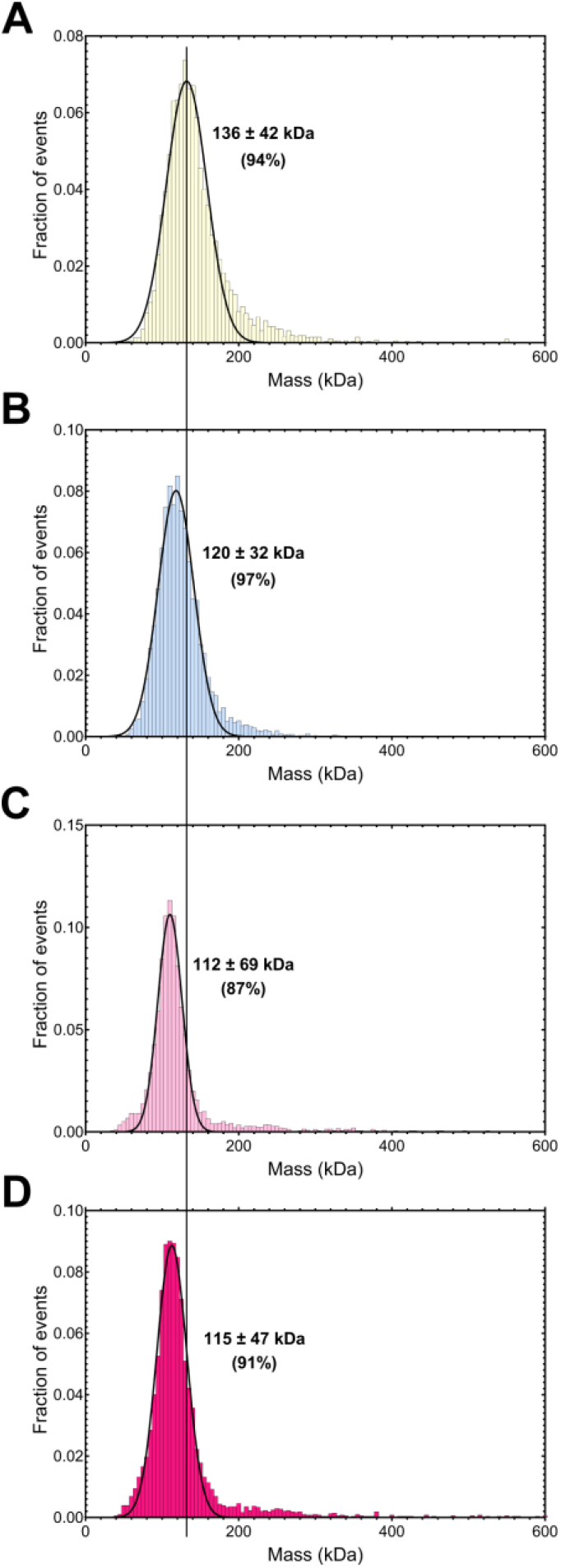
The Rad18 homodimer engages Rad6 in a tight complex. Estimated molecular weight distributions of Rad6(Rad18)_2_ dilutions. Mass photometry histograms were each fit to a single Gaussian function. The kDa value in each plot corresponds to the respective mass at the center of each peak. The percentage indicated for each peak reflects the fraction of events defined by the respective Gaussian function. Panels **A** – **D** show Rad6(Rad18)_2_ complexes diluted to 62.5, 31.25, 12.5, and 6.25 nM, respectively. The vertical line through all plots highlights the shift from a higher molecular weight complex to a lower molecular weight complex as a result of dilution.

### Rad18 stimulates turnover of Uba1 during Rad6 charging

To comprehensively and quantitatively investigate the kinetics of Rad6 charging, we used Cyanine5-labeled ubiquitin (Cy5-Ub) to monitor the covalent attachment of ubiquitin to all proteins present. This allowed Uba1 activation and Rad6 charging to be simultaneously monitored under various kinetic assay conditions. First, the chemistry step of Rad6 charging, i.e., ubiquitin transthioesterification, was investigated by performing kinetic assays under single-turnover conditions with activated Uba1 in excess of substrate, either as free Rad6 or Rad6(Rad18)_2_. Here, activated Uba1 first reversibly engages Rad6, forming the activated Uba1•Rad6 complex (*Step 1*, **Scheme 1**). Next, ubiquitin is transferred from activated Uba1 to Rad6 via ubiquitin transthioesterification, forming the apo Uba1•charged Rad6 complex (*Step 2*, **Scheme 1**). For all assays carried out under this kinetic condition, the observed increases in the concentration of charged Rad6 over time (**Figure 3A–B**) each display a single phase and are best fit to a single exponential rise, yielding observed rate constants for Rad6 charging (*k*_obs,inc_).

**Figure 3.**
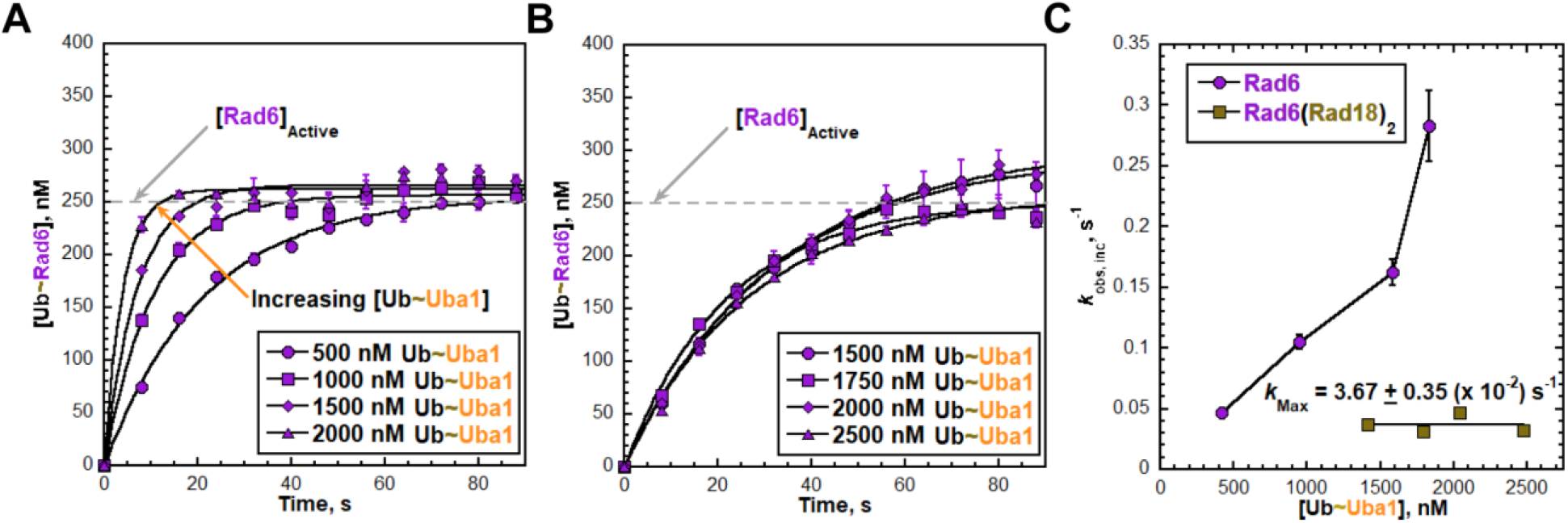
The Rad18 homodimer limits the chemistry step of Rad6 charging by Uba1. Rad6 charging was monitored under single-turnover conditions as described in the **Experimental Procedures. (A - B**) Quantitative analyses. For each concentration of Ub∼Uba1, the concentration of charged Rad6 (Ub∼Rad6 in nM) is plotted as a function of time and all respective data points are fit to a single exponential rise, yielding an observed rate constant, *k*_obs,inc_. Data for each concentration of Ub∼Uba1 represent the average ± S.E.M. of at least three independent experiments. Results from single-turnover assays with free Rad6 and Rad6(Rad18)_2_ are displayed in panels **A** and **B**, respectively. (**C**) Kinetic summary. Rate constants (*k*_obs,inc_) from the data presented in panels **A** (Rad6) and **B** (Rad6(Rad18)_2_) are plotted as a function of the concentration of Ub∼Uba1 measured for each condition (**Figure S5**). Data points for free Rad6 are connected via an interpolated fit for visualization only. Data points for Rad6(Rad18)_2_ are fit to a flat line, yielding a maximal rate constant (*k*_max_).

Furthermore, the initial concentrations of activated Uba1 (i.e., pre-activated Uba1 at t = 0, 500– 2500 nM) are maintained throughout the entirety of all time courses (**Figure S5**), indicating that re-activation of apo Uba1 is significantly faster than the release of charged Rad6 from apo Uba1 (i.e., product release), such that it is essentially instantaneous. Collectively, this indicates that the observed rate constants report on all kinetic steps up to and including the chemistry step (*Step 2*, **Scheme 1**) and are not influenced by re-activation of apo Uba1, product release, or product inhibition.

**Scheme 1.**
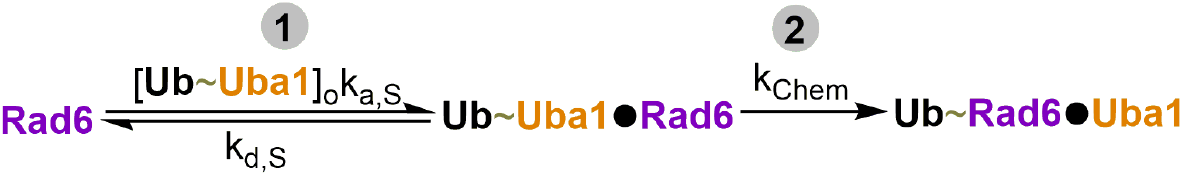
Minimal kinetic scheme for single-turnover assays monitoring Rad6 charging. Step 1: Substrate binding. Step 2: Chemistry.

In assays performed with free Rad6, charging becomes increasingly faster with each successive concentration of activated Uba1 (**Figure 3A**). This indicates that Rad6 charging is rate-limited by substrate binding under these conditions^28^. In assays performed with Rad6(Rad18)_2_, the speed of Rad6 charging appears to be independent of the concentration of activated Uba1 over the examined range (**Figure 3B**), suggesting that all reactions occur with similar rate constants under these conditions. To gain further insights, the observed rate constants from the single exponential fits of the data for Rad6 (**Figure 3A**) and Rad6(Rad18)_2_ (**Figure 3B**) are each plotted in **Figure 3C** as a function of the corresponding concentrations of activated Uba1 (from **Figure S5**). Indeed, *k*_obs,inc_ from assays performed with free Rad6 increases over the entire range of concentrations examined. Under these sub-saturating enzyme conditions, *k*_obs,inc_ is rate-limited by substrate binding and, hence, less than the maximal rate constant (i.e., *k*_chem_) at which charging can proceed. It should be noted that further increases in the concentration of activated Uba1 are not possible because the resulting reactions proceed so quickly that useful time points cannot be collected manually. Thus, only a lower limit for *k*_chem_ can be estimated, such that *k*_chem_ is greater than the largest rate constant observed for charging of free Rad6 (i.e., *k*_chem_ > 0.283 ± 0.029 s^-1^). In contrast, *k*_obs,inc_ from assays performed with Rad6(Rad18)_2_ remains constant over the entire range of activated Uba1 concentrations examined and the observed value (*k*_max_ = 3.67 × 10^-2^ s^-1^ ± 0.35 × 10^-2^ s^-1^) is nearly 8-fold (7.71 ± 1.08-fold) slower than the lower limit for *k*_chem_ (*k*_chem_ > 0.283 ± 0.029 s^-1^) established from assays carried out with free Rad6.

These opposing kinetic behaviors re-affirm the composition of Rad6(Rad18)_2_ utilized in the kinetic assays (i.e., all Rad6 is engaged by Rad18 homodimers) and indicate that *k*_max_ observed for Rad6(Rad18)_2_ reports on a kinetic step, discussed further below, that precedes substrate binding (i.e., *Step 1* in **Scheme 1**). Altogether, the results from these single-turnover kinetic assays (**Figure 3**) reveal that the Rad18 homodimer limits substrate binding and, consequently, the chemistry step of Rad6 charging by Uba1.

Following the chemistry step, apo Uba1 must release the product (charged Rad6) and re-activate to complete its catalytic cycle. To investigate kinetic steps after chemistry, kinetic assays were performed under pre-steady state (i.e., “burst”) conditions with pre-activated Uba1 at a low concentration (2.5 nM) and substrate (either as free Rad6 or Rad6(Rad18)_2_) at an 80- to 320-fold excess. Here, substrates are converted to products through multiple rounds, i.e., turnovers, of enzyme activity in which initial turnovers differ from subsequent turnovers in the kinetic steps involved (depicted in **Scheme 2**). Initial turnovers start with pre-activated Uba1 and Rad6, involve only product formation (which encompasses *Steps 1* and *2*, **Scheme 2**), and convert pre-activated Uba1 and Rad6 to apo Uba1 and charged Rad6. For subsequent turnovers to occur, apo Uba1 must first complete its catalytic cycle by releasing the product (*Step 3*, **Scheme 2**) and re-activating (*Step 4*, **Scheme 2**) and then proceed through product formation (*Steps 1* and *2*, **Scheme 2**). Accordingly, if product release (*Step 3*, **Scheme 2**) and/or re-activation of apo Uba1 (*Step 4*, **Scheme 2**) are slower than product formation (*Steps 1* and *2*, **Scheme 2**), initial turnovers will be faster than all subsequent turnovers. Consequently, product resulting from initial turnovers will accumulate, and a rapid burst of Rad6 charging will be observed, followed by a slower, linear (i.e., steady state) phase^28^.

In each assay performed with free Rad6, the respective data for Rad6 charging after t = 0 conform to a linear regression with a y-intercept that is significantly elevated above the origin (**Figure 4A**, *Top* and *Bottom*). This indicates that a rapid, initial burst of Rad6 charging occurs within the first time point and is followed by a slower, steady state phase. The slope of each linear regression represents the initial velocity (*v*_ss_) of the respective steady state phase.

**Figure 4.**
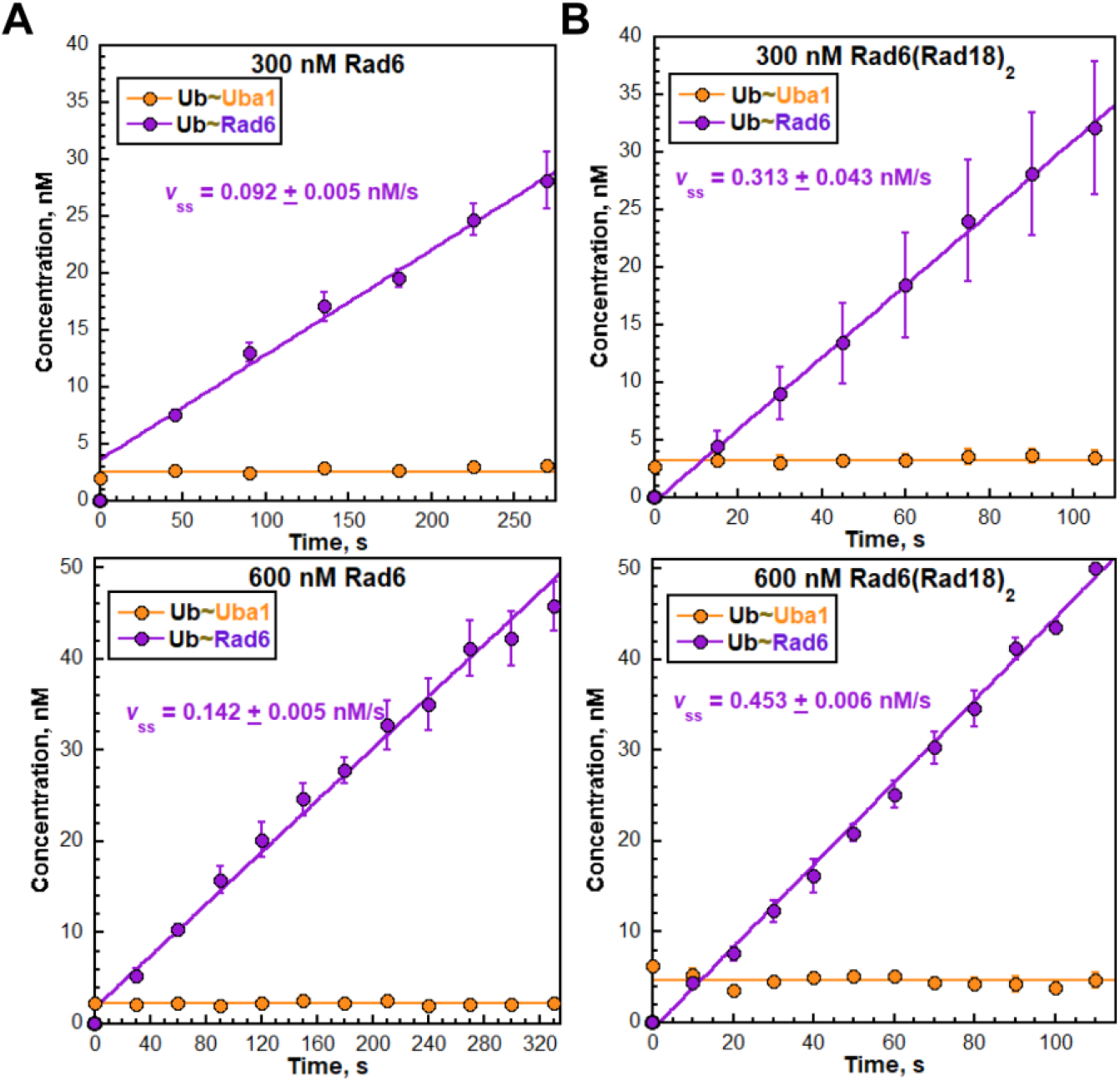
The Rad18 homodimer shifts the rate-limiting step of Rad6 charging by Uba1. Rad6 charging was monitored under pre-steady state (burst) conditions, as described in the **Experimental Procedures. (A – B**) Quantitative analyses. For each condition, the concentration of activated Uba1 (Ub∼Uba1) and charged Rad6 (Ub∼Rad6) are plotted as a function of time. For the former (Ub∼Uba1), all respective data points are fit to a flat line. For the latter (Ub∼Rad6), all respective data points after time zero (i.e., t = 0) are fit to a linear regression yielding a y-intercept that provides an initial estimate of the amplitude of an observed burst and a slope that reflects the initial velocity (*v*_ss_, indicated) of the steady state phase. Data for each condition represent the average ± S.E.M. of at least three independent experiments. Results from assays with free Rad6 and Rad6(Rad18)_2_ are displayed in panels **A** and **B**, respectively. For each panel, results for assays carried out with 300 and 600 nM substrate are displayed in the *Top* and *Bottom*, respectively.

Additionally, the initial concentration of activated Uba1 used in each assay (i.e., pre-activated Uba1 at t = 0) is maintained throughout the entirety of the time course (**Figure 4A**), indicating that apo Uba1 is transient. Accordingly, Uba1 re-activation (*Step 4*, **Scheme 2**) is much faster than, and hence rate-limited by, product release (*Step 3*, **Scheme 2**). This confirms that product release is indeed irreversible under these kinetic conditions. The behaviors observed in **Figure 4A** are reproduced in independent experiments carried out at five different concentrations of Rad6 (**Figure S6**). Altogether, this indicates that product release (*Step 3*, **Scheme 2**) is significantly slower than all other kinetic steps and, hence, is rate-limiting with free Rad6 under these kinetic conditions. Thus, Uba1 retains significant affinity for Rad6 following charging. In other words, the affinity of activated Uba1 for the Rad6 substrate is at least comparable to the affinity of apo Uba1 for the charged Rad6 product. This agrees with results from a prior study of human Rad6 and Uba1^27^ that suggested the affinity of Uba1 for free Rad6 is not significantly attenuated by the processes of Uba1 activation and/or Rad6 charging, as in the proposed thioester “affinity switch” model^30^.

In each assay performed with Rad6(Rad18)_2_, the initial concentration of activated Uba1 (i.e., pre-activated Uba1 at t = 0) is maintained throughout the entirety of the time course (**Figure 4B**), as observed for the corresponding assays with free Rad6 (**Figure 4A, Figure S6**). Thus, Uba1 re-activation (*Step 4*, **Scheme 2**) is much faster than and rate-limited by product release (*Step 3*, **Scheme 2**), product release is irreversible, and these behaviors are independent of the Rad18 homodimer. However, with Rad6(Rad18)_2_, the respective data for Rad6 charging after t = 0 conform to a linear regression that intersects the origin (**Figure 4B**). In other words, a burst is not observed, and the steady state is established instantaneously. This indicates that with Rad6(Rad18)_2_, initial turnovers are significantly slower than subsequent turnovers and, therefore, the rate-limiting step occurs within product formation (*Steps 1* and *2*, **Scheme 2**), in contrast to the behavior observed with free Rad6 (**Figure 4A**)^28^. Collectively, these results indicate that interactions of the Rad18 homodimer with Rad6 shift the rate-limiting step from product release (*Step 3*, **Scheme 2**) to a step within product formation (*Steps 1* and *2*, **Scheme 2**).

The opposing behaviors of activated Uba1 with free Rad6 and Rad6(Rad18)_2_ observed in **Figure 4** indicate that interactions of the Rad18 homodimer with Rad6 either; 1) decrease the speed of initial turnovers; 2) increase the speed of subsequent turnovers, or; 3) cause both effects^28^. The results from single-turnover kinetic assays presented in **Figure 3** reveal that interactions of the Rad18 homodimer with Rad6 limit substrate binding (*Step 1*, **Scheme 2**) and, consequently, chemistry (*Step 2*, **Scheme 2**). Accordingly, the speed of initial turnovers (*Steps 1* and *2*, **Scheme 2**) decreases. If this were solely responsible for the elimination of the burst phases in pre-steady state kinetic assays carried out with Rad6(Rad18)_2_ (**Figure 4B**), then the corresponding initial velocities would also be slower than those observed in assays carried out with free Rad6 (**Figure 4A**)^28^. However, for each concentration of substrate, the initial velocities are significantly faster with Rad6(Rad18)_2_ than with free Rad6. Specifically, at 300 nM substrate, the initial velocity with Rad6(Rad18)_2_ is 3.40 ± 0.50-fold faster (**Figure 4B**, *Top*) than that observed with free Rad6 (**Figure 4A**, *Top*). At 600 nM substrate, the initial velocity with Rad6(Rad18)_2_ is 3.19 ± 0.13-fold faster (**Figure 4B**, *Bottom*) than that observed with free Rad6 (**Figure 4A**, *Bottom*). This suggests that, in addition to slowing initial turnovers by limiting substrate binding (*Step 1*, **Scheme 2**), interactions of the Rad18 homodimer with Rad6 also significantly stimulate product release (*Step 3*, **Scheme 2**) such that Uba1 turnover is ultimately accelerated beyond that observed with free Rad6. To confirm this, kinetic assays were performed under steady state conditions with pre-activated Uba1 at a very low concentration (0.25 nM) and substrate (either as free Rad6 or Rad6(Rad18)_2_) at a very high excess (800-fold).

In steady state kinetic assays, substrates are converted to products through multiple turnovers of enzyme activity, similar to the pre-steady state assays described above, except that initial turnovers are negligible due to the very low concentration of pre-activated enzyme^28^. Hence, essentially all Rad6 charging occurs via subsequent turnovers (depicted in **Scheme 3**), which each start with product release, proceed instantaneously through Uba1 re-activation, and are completed by product formation. In all assays, the initial concentrations of activated Uba1 (i.e., pre-activated Uba1 at t = 0) are maintained throughout the time courses (**Figure 5**), as expected. Again, this re-affirms that Uba1 re-activation (*Step 2*, **Scheme 3**) is much faster than and rate-limited by product release (*Step 1*, **Scheme 3**), product release is irreversible, and these behaviors are independent of the Rad18 homodimer. For assays performed with free Rad6, *v*_ss_ = 0.256 ± 0.012 nM/s (**Figure 5A**). The results presented in **Figure 4A** for free Rad6 indicate that product release (*Step 1*, **Scheme 3**) is significantly slower than all subsequent steps (Steps 2 – 4, **Scheme 3**) and, therefore, is the rate-limiting step. Hence, the initial velocity observed with free Rad6 in **Figure 5A** reflects product release. The results presented in **Figure 4B** for Rad6(Rad18)_2_ indicate that substrate binding (*Step 3*, **Scheme 3**) is significantly slower than all other kinetic steps (*Steps 1, 2*, and *4*, **Scheme 3**) and, therefore, is rate-limiting. Hence, the initial velocity observed with Rad6(Rad18)_2_ (**Figure 5B**) reflects substrate binding. For assays performed with Rad6(Rad18)_2_, the initial velocity (*v*_ss_ = 0.708 ± 0.005 nM/s, **Figure 5B**) is 2.76

**Figure 5.**
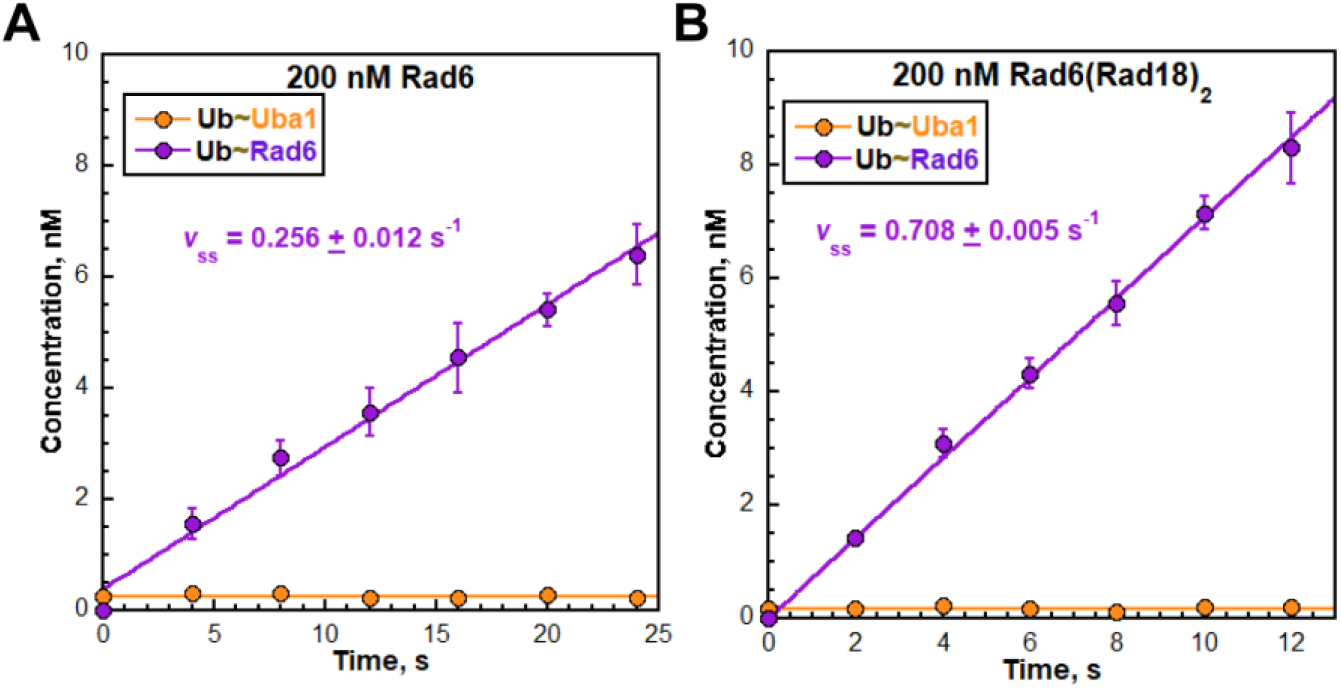
The Rad18 homodimer stimulates the product release step of Rad6 charging by Uba1. Rad6 charging was monitored under steady state conditions, as described in the **Experimental Procedures. (A – B**) Quantitative analyses. For each condition, the concentration of activated Uba1 (Ub∼Uba1) and Rad6 ubiquitin thioester conjugates (Ub∼Rad6) are plotted and analyzed as in **Figure 4** above. Data for each condition represent the average ± S.E.M. of at least three independent experiments. Results from assays with free Rad6 and Rad6(Rad18)_2_ are displayed in panels **A** and **B**, respectively.

**Scheme 3.**
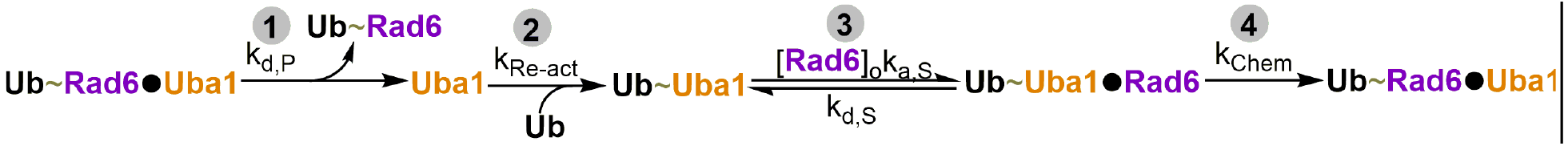
Minimal kinetic scheme for steady state assays monitoring Rad6 charging. Step 1: Product release. Step 2: Re-activation of apo Uba1. Step 3: Substrate binding. Step 4: Chemistry.

± 0.13-fold faster than observed with free Rad6 (*v*_ss_ = 0.256 ± 0.012 nM/s, **Figure 5A**). This is only possible if all kinetic steps up to and including substrate binding (*Steps 1* – *3*, **Scheme 3**) with Rad6(Rad18)_2_ are faster than product release with free Rad6 (*Step 1*, **Scheme 3**) under these kinetic assay conditions. These results confirm the hypothesis described above that interactions of the Rad18 homodimer with Rad6 stimulate product release, thereby accelerating Uba1 turnover beyond that observed with free Rad6.

**Scheme 2.**
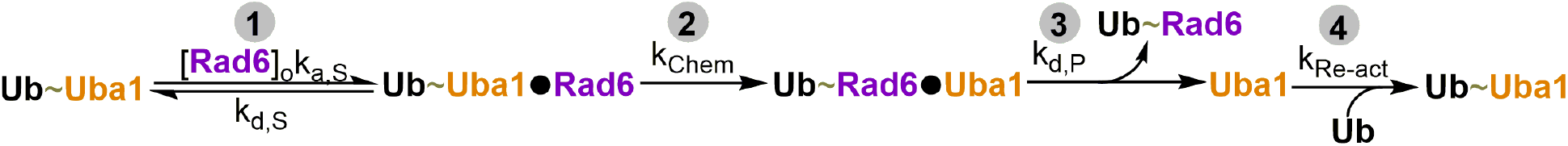
Minimal kinetic scheme for pre-steady state (“burst”) assays monitoring Rad6 charging. Step 1: Substrate binding. Step 2: Chemistry. Step 3: Product release. Step 4: Re-activation of apo Uba1. The order of kinetic steps for initial turnovers is 1→2. The order of kinetic steps for subsequent turnovers is 3→4→1→2.

### Rad6 and charged Rad6 are engaged by Rad18 homodimers with identical affinities

Charging of free Rad6 in the nuclear pool of this E2 must ultimately be accompanied by binding of a Rad18 homodimer. Currently, it is unclear how this occurs in the context of the nucleus, where charged Rad6 is competing with Rad6 for interaction with Rad18 homodimers (**Scheme 4**). To investigate this, we utilized isothermal titration calorimetry (ITC) to measure and directly compare the affinities, in the form of dissociation constants (K_D_), of Rad18 homodimers for Rad6 and a charged Rad6 mimic (Rad6^iso^-Ub). Specifically, homodimers of SUMO-Rad18 were titrated with either native Rad6 or Rad6^iso^-Ub in which the active-site cysteine (C88) is mutated to lysine and covalently attached to the C-terminal glycine of ubiquitin by a stable isopeptide bond^24,25^. Importantly, the stoichiometries of the interactions analyzed by ITC (Rad6:SUMO-Rad18 = 0.513 ± 0.005, Rad6^iso^-Ub:SUMO-Rad18 = 0.505 ± 0.005) agree exceptionally well with the stoichiometry of the native Rad6(Rad18)_2_ complex (**Figure 6**). The tight affinity of (SUMO-Rad18)_2_ for Rad6 (91.515 ± 10.328 nM, **Figure 6**) determined by ITC agrees with the tight affinity of native Rad6(Rad18)_2_ observed by mass photometry (**Figure 2**) and other techniques in our lab^13^. Thus, the His-SUMO tag on Rad18 does not significantly affect the affinity of the Rad18 homodimer for Rad6. The affinity of (SUMO-Rad18)_2_ for Rad6^iso^-Ub (85.505 ± 10.371 nM, **Figure 6**) is within experimental error of that observed for Rad6 (91.515 ± 10.328 nM) and the difference between the affinities (ΔK_D_ = 6.010 ± 14.636 nM) is not statistically significant (two-tailed p-value = 0.7024). The identical affinities observed by ITC indicate that binding of (SUMO-Rad18)_2_ to either Rad6 form is not thermodynamically favored over the other. This suggests that the interactions between Rad6 and Rad18 homodimers are independent of the ubiquitin thioester status of Rad6 and, hence, not directed by a thioster “affinity switch,” in contrast to that suggested for other RING E3’s^25,31-34^ This has significant implications for PCNA monoubiquitination and RING E3-dependent ubiquitination cascades in general, as discussed further below.

**Figure 6.**
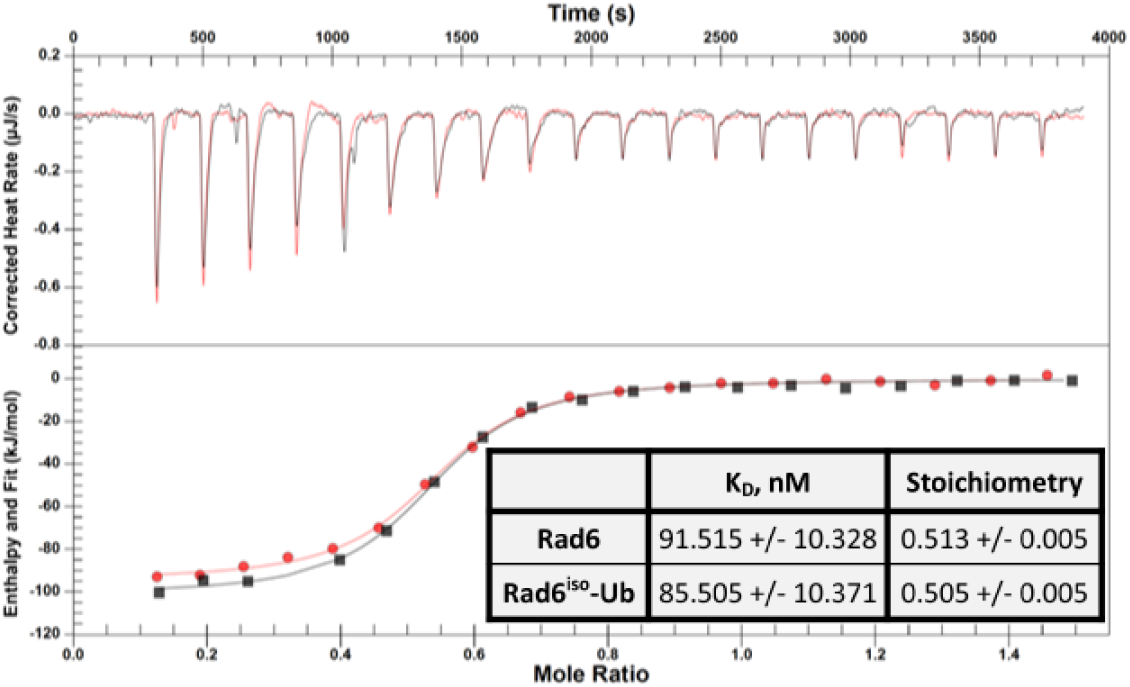
The Rad18 homodimer engages Rad6 and charged Rad6 with identical affinities. Binding of (SUMO-Rad18)_2_ to Rad6 isoforms was monitored by ITC, as described in the **Experimental Procedures**. Data from representative, paired titrations with Rad6 and Rad6^iso^-Ub are displayed in black and red, respectively, and overlaid for direct comparison. Heat rate peaks are displayed in the top panel as a function of time. The corresponding binding isotherms are displayed in the bottom panel as a function of the molar ratios resulting from the injections and each are fit to an independent one-site binding model yielding a dissociation constant (K_D_) and stoichiometry. The average (± S.E.M.) K_D_ and stoichiometry values for each Rad6 isoform from three independently paired titrations are reported in the inset table.

**Scheme 4.**
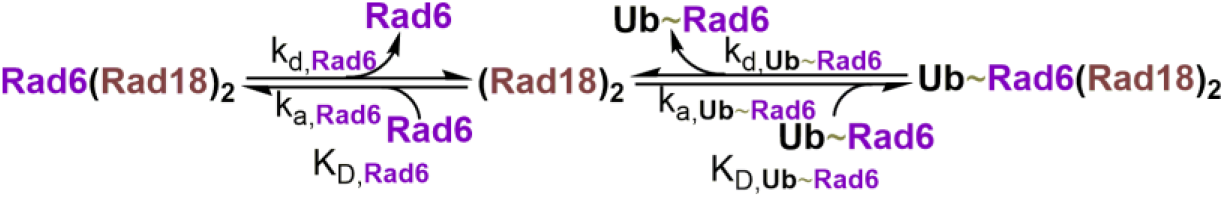
Competition between Rad6 and charged Rad6 for binding to the Rad18 homodimer.

## Discussion

In the present study, we utilize human Rad6(Rad18)_2_ as a model bivalent E2:RING E3 complex to delineate the interplay of protein•protein interactions among Uba1, an E2, and a RING E3 during E2 charging. Specifically, we monitored Uba1 activation and Rad6 charging simultaneously via fluorescence-based kinetic assays (**Figure 1**). This permitted the charging of free Rad6 to be directly compared to the charging of Rad6(Rad18)_2_ complexes across conditions that highlight distinct mechanistic steps of the Uba1 catalytic cycle (**Figures 3 –** 5). Mass photometry analyses (**Figure 2**) established that, prior to incubation with activated Uba1, Rad6(Rad18)_2_ exists exclusively as a tight complex at the concentrations utilized in all kinetic assays of the current study (i.e., all Rad6 is engaged by Rad18 homodimers with high affinity). Results from single-turnover kinetic assays reveal that interactions of the Rad18 homodimer with Rad6 slow the binding of activated Uba1 and, consequently, the chemistry step of Rad6 charging (**Figure 3**). Thus, *k*_max_ observed for Rad6(Rad18)_2_ under these kinetic assay conditions reports on a mechanistic step that precedes activated Uba1 binding to Rad6. As alluded to above, Rad6 in complex with a Rad18 homodimer must disengage from (at least) a RING domain of (Rad18)_2_ in order to be engaged by activated Uba1 for subsequent charging ^1,4-6^. Correspondingly, interactions of Rad6 with a RING domain of (Rad18)_2_ are expected to limit binding of activated Uba1 via competitive inhibition and, consequently, the chemistry step of Rad6 charging^35^. This suggests that *k*_max_ observed for Rad6(Rad18)_2_ in **Figure 3** reports on dissociation of Rad6 from (at least) a RING domain within the Rad18 homodimer. Results from pre-steady state (**Figure 4, Figure S6**) and steady state (**Figure 5**) kinetic assays reveal that interactions of the Rad18 homodimer with charged Rad6 significantly stimulate release of the charged Rad6 product from apo Uba1 such that Uba1 turnover is ultimately accelerated beyond that observed with free Rad6. Consequently, more Rad6 is charged by Uba1 per unit time when Rad6 is pre-engaged by (Rad18)_2_ in a complex compared to when Rad6 is free in solution. In other words, Rad6 engaged by a Rad18 homodimer is a better substrate than free Rad6 when Uba1 is limiting, which is likely the scenario *in vivo* where cellular E2’s are present at > 400-fold excess of Uba1^36^.

Collectively, the results from the comprehensive kinetic assays presented in the current study reveal a novel mechanism for the charging of a bivalent E2:RING E3 complex by Uba1 in which the respective RING E3 slows certain kinetic steps of the Uba1 catalytic cycle (i.e., substrate binding, chemistry) while accelerating others (product release), ultimately stimulating E2 charging overall. To the best of our knowledge, this represents the first example of a RING E3 stimulating the Uba1-dependent charging of an E2. The observed stimulation may occur by one of at least two mechanisms that are distinguished by the nature of the interactions between Rad18 homodimers and Rad6. In one pathway, referred to as macroscopic dissociation, the Uba1-binding site of Rad6 is exposed to activated Uba1 via complete release of the E2 substrate from all interactions with a Rad18 homodimer (described by the rate constant, *k*_max_, Figure 3C). After charging, the process of the Rad18 homodimer forming interactions with charged Rad6 stimulates release of the charged E2 product from apo Uba1 (described by the rates in **Figures 4B** and **5B**). In this pathway, macroscopic release of Rad6 from a Rad18 homodimer renders the subsequent formation of charged Rad6(Rad18)_2_ complexes susceptible to inhibition by any free Rad6 in the nuclear pool. The results from the ITC experiments presented in **Figure** 6 above suggest that the interactions of Rad18 homodimers with the substrate (Rad6) and product (charged Rad6) of the Uba1 catalytic cycle are not directed by a thioester “affinity switch”.

Consequently, this pathway requires the population of free, charged Rad6 within the nuclear Rad6 pool to be in significant excess of “discharged” Rad6 in order to effectively compete for binding to Rad18 homodimers, forming charged Rad6(Rad18)_2_ complexes, and ultimately monoubiquitinate PCNA. In an alternative pathway, referred to as microscopic dissociation, a R6B domain within a Rad18 homodimer maintains constant contact with the “backside” of Rad6 while the Uba1-binding site of Rad6 is released from a Rad18 RING domain (described by the rate constant, *k*_max_, **Figure 3C**) and subsequently engaged by activated Uba1. After charging, the process by which the resident Rad6 is re-engaged by a Rad18 RING domain stimulates release of the charged E2 product from apo Uba1 (described by the rates in **Figures 4B** and **5B**). In this pathway, the propensity for Rad6 to remain engaged to a Rad18 homodimer (via a R6B domain) throughout the Uba1 catalytic cycle and the lack of a thioester “affinity switch” directing interactions between Rad6 and Rad18 homodimers (noted above) collectively diminish the capacity of free Rad6 in the nuclear pool to inhibit formation of charged Rad6(Rad18)_2_ complexes.

In conclusion, results from the thermodynamic, biophysical, and comprehensive kinetic studies presented in the current study reveal an overall novel mechanism of charging a bivalent E2:RING E3 complex that is not directed by ubiquitin thioester “affinity switches.” Rather, (Rad18)_2_ stimulates Rad6 charging by accelerating release of the Ub∼Rad6 product from apo Uba1. Currently, studies are underway to decipher which of the pathways described above for Rad18-dependent stimulation of Rad6(Rad18)_2_ charging by Uba1 are valid and whether the RING E3-dependent charging observed in the present study for Rad6(Rad18)_2_ is conserved in other bivalent E2:RING E3 complexes.

## Supporting information

Supplemental Experimental Procedures, Results, and Figures

## SUPPLEMENTAL INFORMATION

Supplemental Information includes Supplemental Experimental Procedures, Supplemental Results, Supplemental Figures S1 – S6, and can be found with this article online at…..

## ACKNOWLEDGEMENTS

The authors thank Dr. Neela Yennawar and Ms. Julia Fecko for their assistance and helpful discussions related to the ITC binding studies. ITC measurements were performed using an instrument housed in the Automated Biological Calorimetry Core Facility at Penn State’s Huck Institutes of the Life Sciences, which was supported by NIH grant S10OD025145 to Dr. Yennawar. This work was supported by funding from the National Institutes of Health to M.H. (R35 GM147238) and E.A. (R01 GM130756, R01 GM133967, R35 GM149320, and S10 OD030343).

## AUTHOR CONTRIBUTIONS

S.P. modified, expressed, and purified all proteins. M.H., S.P., A.M., and E.A. designed the experiments. S.P. and A.M. performed the experiments. M.H., S.P., A.M., and E.A. analyzed the data and wrote the paper.

