## Supplemental Experimental Procedures, Results, and Figures for "A novel mechanism of ubiquitin-charging a bivalent E2:RING E3 complex"

**Affiliations**:
[1]Department of Chemistry, The Pennsylvania State University, University Park, PA 16802
[2]Department of Biochemistry & Molecular Biology, Saint Louis University School of Medicine, Saint Louis, MO 63104


### Supplemental Experimental Procedures

*Determining the concentration of active Uba1*

All concentrations indicated below are final (i.e., after mixing). The stoichiometric formation of Uba1 ubiquitin thioester conjugates was monitored using a modified protocol (**Figure S2A**)[1-5]. First, ATP (1 mM) was pre-incubated with a 1X ATP regeneration system (10 mM phosphocreatine, 10 units/mL creatine phosphokinase) to generate a uniform ATP solution. Next, Cy5-Ub (5 μM) was added, and the reaction was initiated by the addition of Uba1 ($\geq$ 500 nM absolute Uba1). Aliquots were removed every 10 s and quenched into 1X non-reducing loading buffer. After all time points were completed, samples were resolved via gel electrophoresis, the gels were imaged (**Figure S2B**) and quantitatively analyzed (**Figure S2C**) as described in the main text. For all Uba1 samples used in the present study, the concentration of active Uba1 was typically ~50% of the absolute concentration of Uba1.

*Determining the concentration of active Rad6*

All concentrations indicated below are final (i.e., after mixing). The stoichiometric formation of Rad6 ubiquitin thioester conjugates via Uba1-catalyzed ubiquitin transthioesterification was monitored using a modified protocol (**Figure S3A**)[6]. First, ATP (1 mM) was pre-incubated with a 1X ATP regeneration system to generate a uniform ATP solution. Next, Uba1 (600 nM active protein) and Cy5-Ub (5 μM) were added in succession, and the resultant solution was pre-incubated to form stoichiometric Uba1~ubiquitin thioester conjugates. Finally, the reaction was initiated by the addition of Rad6 (300 nM absolute Rad6, either as free Rad6 or Rad6 within $Rad6(Rad18)_2$). Aliquots (2 μL) were removed at the indicated time points and quenched into 58 μL of pre-diluted loading buffer to achieve a 30-fold dilution and final 1X quench composition (50 mM Tris•HCl, 10% glycerol, 2% SDS, 175 mM EDTA). After all time points were completed, samples were resolved via gel electrophoresis, and the gels were imaged (**Figure S3B**) and quantitatively analyzed (**Figure S3C**) as described in the main text. All Rad6 samples utilized in the present study were 100% active (i.e., the concentrations of active Rad6 calculated from the assays are equal to the absolute concentrations of Rad6 utilized in the assays).

### Supplemental Results

*Human Rad6(Rad18)$_2$ is predicted to be highly stable*

To estimate the stability of the human Rad6(Rad18)$_2$ complex, we used an additive binding energy model[7] that estimates the overall dissociation constant of the complex, $K_{D,Ov}$, based on the individual and independent affinities of Rad6 for an isolated Rad18 RING domain and an isolated Rad18 R6B domain. Previous studies demonstrated that interactions of Rad6 with a Rad18 RING domain and a Rad18 R6B domain are independent of each other (reviewed in [8]). The model posits that the negative common logarithm of the overall dissociation constant, $K_{D,Ov}$, of the complex (i.e., $-\log_{10}K_{D,Ov}$) is, on average, one log unit less than the sum of the negative common logarithms of the dissociation constants of the individual, independent binding domains (i.e., $-\log_{10}K_{D,R6B} + -\log_{10}K_{D,RING}$) (**Equation 1**).

$$\mathbf{1}\quad -\log_{10}K_{D,Ov} = -\log_{10}K_{D,R6B} + -\log_{10}K_{D,RING} - 1$$

**Equation 1** can be rewritten as **Equations 2 – 5** and then simplified to **Equation 6** to solve for $K_{D,Ov}$.

$$\mathbf{2}\quad -\log_{10}K_{D,Ov} = -\log_{10}K_{D,R6B} + -\log_{10}K_{D,RING} + \log_{10}0.1$$

$$\mathbf{3}\quad \log_{10}0.1 + \log_{10}K_{D,Ov} = \log_{10}K_{D,R6B} + \log_{10}K_{D,RING}$$

$$\mathbf{4}\quad \log_{10}(K_{D,Ov} \times 0.1) = \log_{10}(K_{D,R6B} \times K_{D,RING})$$

$$\mathbf{5}\quad K_{D,Ov} \times 0.1 = K_{D,R6B} \times K_{D,RING}$$

$$\mathbf{6}\quad K_{D,Ov} = K_{D,R6B} \times K_{D,RING} \times 10$$

The affinity of the isolated Rad18 RING domain for Rad6 was measured by isothermal titration calorimetry to be 35 μM[9]. The affinity of the isolated Rad18 R6B domain for Rad6 was measured by surface plasmon resonance to be 62.1 $\pm$ 8.9 μM and confirmed by NMR[10]. Substituting these values into **Equation 6** yields a value of ~22 nM for $K_{D,Ov}$, as described below, suggesting that the interaction of Rad6 with a Rad18 homodimer is very tight and the resulting complex is highly stable.

$$K_{D,Ov} = (35 \times 10^{-6}\ \text{M}) \times (62.1 \times 10^{-6}\ \text{M}) \times 10 = 21.735 \times 10^{-9}\ \text{M}$$

*Stoichiometric formation of Uba1 ubiquitin thioester conjugates is rapid and maintained*

To determine the concentration of active Uba1, defined here as the Uba1 that can be conjugated to ubiquitin via thioester formation with active-site residue C632, we monitored the stoichiometric formation of Uba1 ubiquitin thioester conjugates using a modified protocol (see **Supplemental Experimental Procedures**)[1-5]. As observed in **Figure S2C**, the concentration of Uba1 ubiquitin thioester conjugates rapidly increases and plateaus within the first time point (10 s). The plateau is maintained for $\geq$ 100 s. Assuming first-order kinetics and that the plateau is reached in $\leq$ 6 half-lives (i.e., 98.43% reaction in 10 s, 10 s = 6 half-lives), the rate constant is estimated to be $\geq 0.416\ \text{s}^{-1}$ ( $k = \frac{\ln(2)}{t_{1/2} \times \frac{10s}{6t_{1/2}}} = 0.416\ s^{-1}$). The concentration of active Uba1 is determined from the observed plateau (i.e., the amplitude for Uba1 ubiquitin thioester formation). As observed in **Figure S2C**, the concentration of active Uba1 used in the assay (194.7 $\pm$ 5.5 nM) is approximately 50% of the concentration of absolute Uba1 (400 nM) utilized in the assay. Hence, the Uba1 preparation utilized in this assay is ~50% active. All Uba1 samples used in the present study were ~30–50% active.

*$(Rad18)_2$ does not affect the amount of Rad6 ubiquitin thioester conjugates*

To determine the concentration of Rad6 that can be conjugated to ubiquitin via Uba1-catalyzed ubiquitin transthioesterification (i.e., active Rad6), we monitored endpoint Rad6 ubiquitin thioester conjugate formation using a modified protocol (see **Supplemental Experimental Procedures**)[6]. Under the single-turnover conditions of the assay (i.e., $[Uba1]_{active}/[Rad6]$ ratios $\geq$ 1.0), the endpoint is stoichiometric with the concentration of active Rad6[6]. As observed in **Figure S3C**, the concentrations of Rad6 ubiquitin thioester conjugates increase to the concentration of absolute Rad6 (300 nM) within the first time point and remain constant for > 20 min. This behavior was observed both in the absence and presence of $(Rad18)_2$. Together, this indicates that 1) the concentration of active Rad6 is equal to the absolute concentration of Rad6 (300 nM) utilized in the assays (i.e., Rad6 is 100% active); 2) Uba1 maintains the maximal concentrations of Rad6 ubiquitin thioester conjugates over a significant period of time; and 3) the Rad18 homodimer does not affect the amount of Rad6 that undergoes Uba1-catalyzed ubiquitin transthioesterification. All Rad6 samples utilized in the present study were 100% active.

**Supplemental Figures**

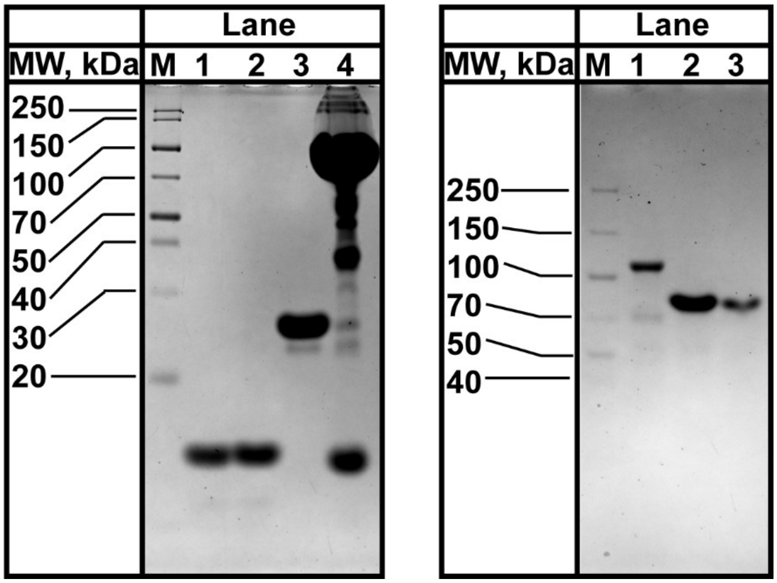


**Figure S1**. Denaturing SDS-PAGE analyses of in-house preparations of recombinant human proteins used in the current study. (*Left*) Marker (Lane M), Rad6 (Lane 1, 300 pmol), $Rad6^{C88K}$ (Lane 2, 300 pmol), $Rad6^{C88K}$-Ub (i.e., $Rad6^{iso}$-Ub, Lane 3, 200 pmol), and $Rad6(Rad18)_2$ (Lane 4, 300 pmol Rad6, 600 pmole Rad18) were loaded on a denaturing 15 % polyacrylamide SDS gel and stained with Coomassie Blue. (*Right*) Marker (Lane M), Uba1 (Lane 1, 20 pmol), $Rad6(Rad18)_2$ (Lane 2, 50 pmol Rad6, 100 pmol Rad18), and SUMO-Rad18 (Lane 3, 50 pmol) were loaded on a Mini-PROTEAN® TGX™ denaturing 4 – 20% gradient precast polyacrylamide gel and stained with Coomassie Blue. The same sample is run in Lanes 4 and 2 of the *Left* and *Right* gels, respectively. Lane 4 of the *Left* gel was overloaded, with respect to Rad18 (600 pmol) to highlight Rad6 (300 pmol). Lane 2 of the *Right* gel was underloaded, with respect to Rad6 (50 pmol, not visible), to highlight Rad18 (100 pmol).

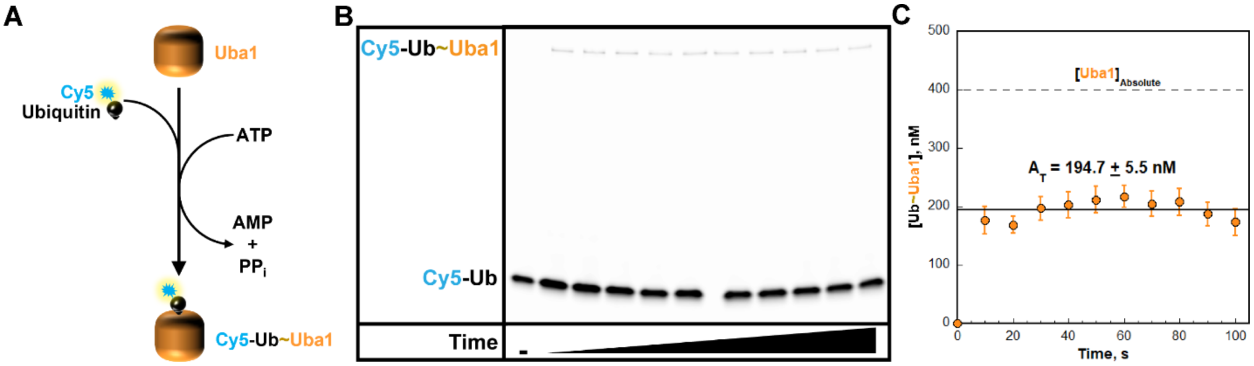


**Figure S2**. Concentration of active Uba1. (**A**) Schematic representation of the fluorescence-based, single-turnover, ubiquitin transfer assay to determine the concentration of active Uba1. Uba1 utilizes ATP to conjugate Cy5-Ub to its active-site cysteine via a high-energy thioester bond. (**B**) Representative fluorescence scan of an assay carried out with 400 nM absolute Uba1. Cy5-labeled ubiquitin (Cy5-Ub) and Uba1 ubiquitin thioester (Cy5-Ub~Uba1) are indicated, where "~" denotes a high-energy thioester bond. (**C**) Quantitative analysis of Uba1 ubiquitin thioester formation. Data represent the average ± S.E.M. of six independent experiments carried out with 400 nM absolute Uba1 (indicated on the plot by a dashed line). The concentration of Uba1 ubiquitin thioester conjugates is plotted as a function of time, and the data points after $t = 0$ are fit to a flat line where the concentration of active Uba1 used in the assays (indicated) is equal to the y-intercept, i.e., amplitude ($A_T$).

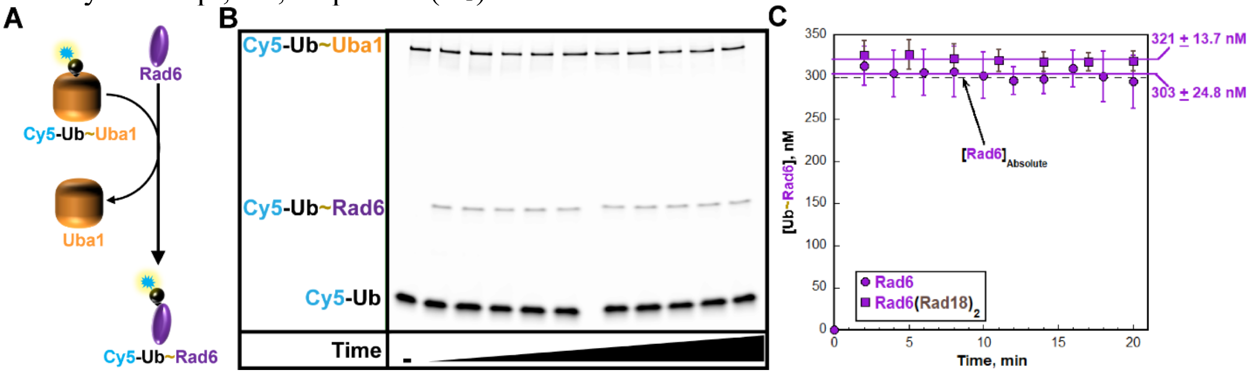


**Figure S3**. Concentration of active Rad6. (**A**) Schematic representation of the fluorescence-based, single-turnover, ubiquitin transfer assay to determine the concentration of active Rad6. Uba1, pre-activated with Cy5-Ub, catalyzes conjugation of Cy5-Ub to Rad6 residue C88 via a high-energy thioester bond. (**B**) Representative fluorescence scan of an assay carried out with 300 nM absolute Rad6. Cy5-labeled ubiquitin (Cy5-Ub), Uba1 ubiquitin thioester (Cy5-Ub~Uba1), and Rad6 ubiquitin thioester (Cy5-Ub~Rad6) are indicated, where "~" denotes a high-energy thioester bond. (**C**) Quantitative analysis of Rad6 ubiquitin thioester formation. The concentration of Rad6 ubiquitin thioester conjugates is plotted as a function of time for assays carried out with either free Rad6 or Rad6 within $Rad6(Rad18)_2$. Data for each time course represent the average ± S.E.M. of at least four independent experiments carried out with 300 nM absolute Rad6 (indicated on the plot by a dashed line), either as free Rad6 or Rad6 within $Rad6(Rad18)_2$. For each time course, data points after $t = 0$ are fit to a flat line where the y-intercept is equal to the concentration of active Rad6 present in the assays (indicated), i.e., amplitude ($A_T$).

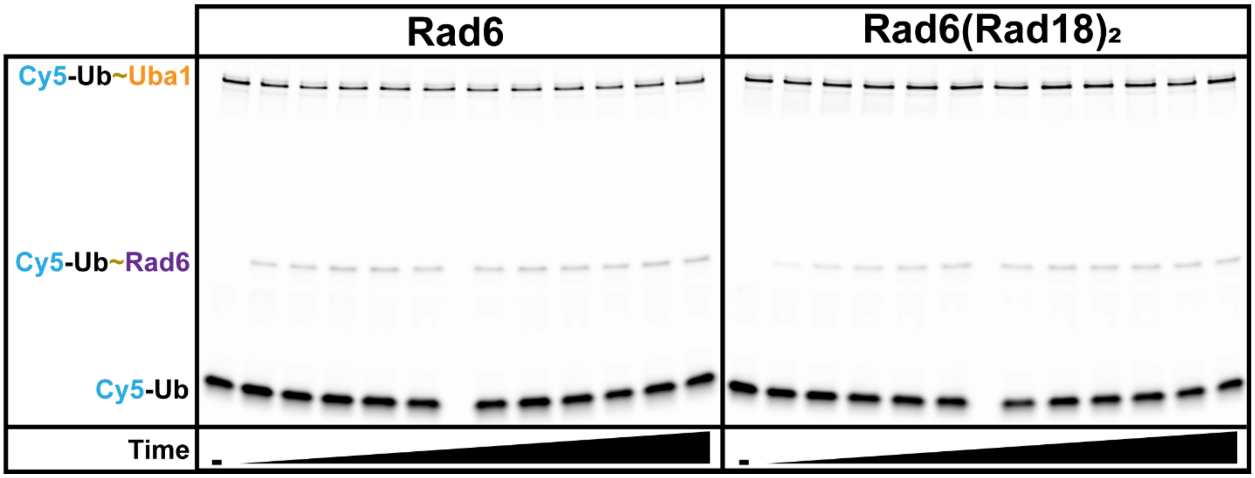


**Figure S4**. Representative fluorescence scan of single-turnover assays carried out with 250 nM active Rad6, either as free Rad6 (*Left*) or Rad6 within Rad6(Rad18)$_2$ (*Right*). Cy5-labeled ubiquitin (Cy5-Ub), Uba1 ubiquitin thioester (Cy5-Ub~Uba1), and Rad6 ubiquitin thioester (Cy5-Ub~Rad6) are indicated, where "~" denotes a high-energy thioester bond.

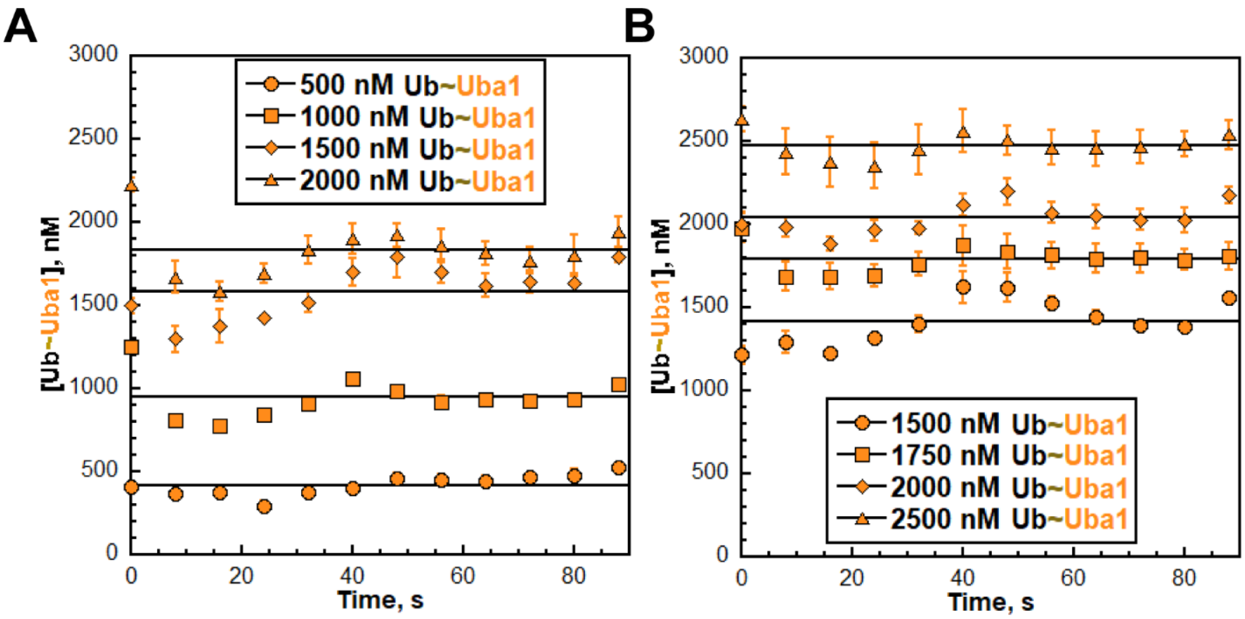


**Figure S5**. The initial concentrations of activated Uba1 are maintained during single-turnover kinetic assays monitoring the formation of Rad6 ubiquitin thioester conjugates. The concentrations of activated Uba1 (i.e., Ub~Uba1, Uba1 ubiquitin thioester conjugates) observed in single-turnover kinetic assays displayed in **Figure 3** of the main text are plotted as a function of time. Panels **A** and **B** show the data observed for assays carried out with free Rad6 (**Figure 3A**) and Rad6(Rad18)$_2$ (**Figure 3B**), respectively. Data points for each condition in both panels are fit to a flat line, yielding a y-intercept that is equivalent to the concentration of Ub~Uba1 that is maintained throughout the respective experiment. The concentrations are used to plot the data in **Figure 3C** in the main text.

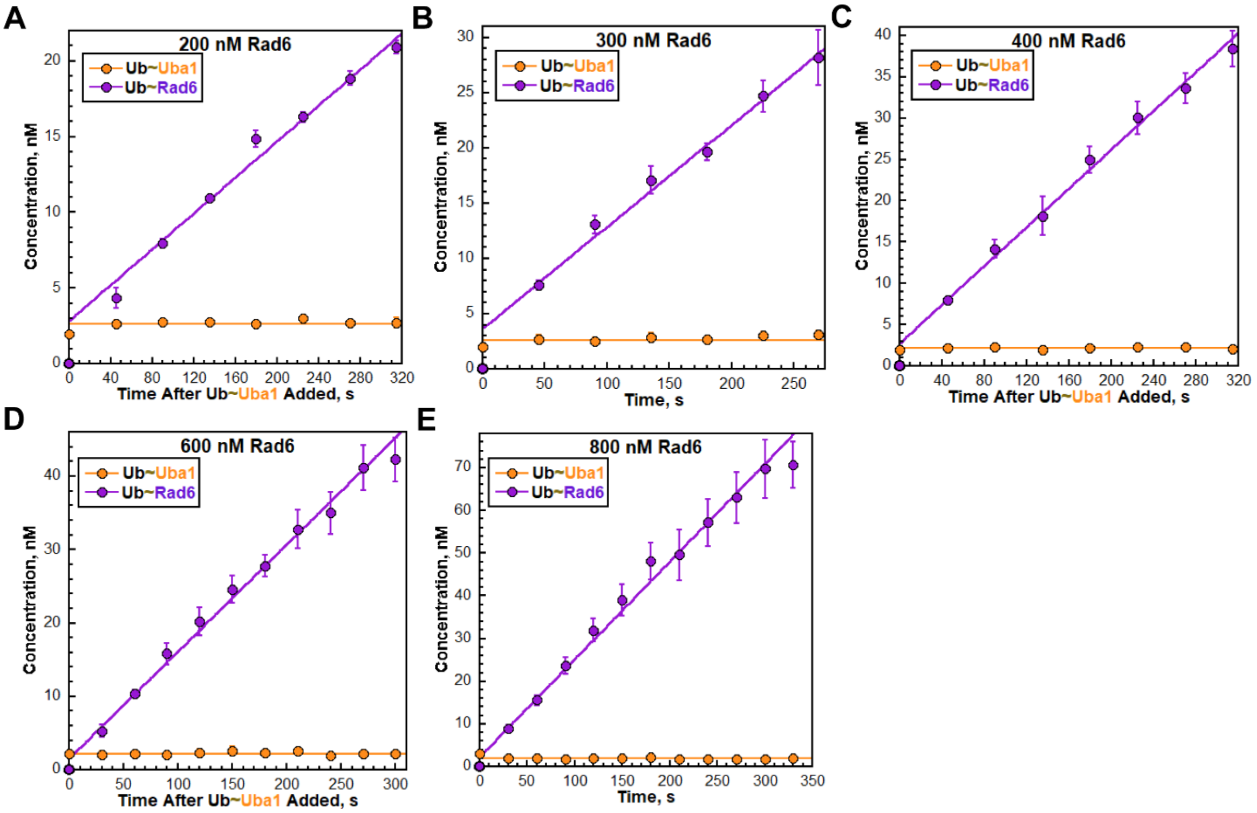


**Figure S6**. A burst of Rad6 charging is observed in pre-steady-state assays with free Rad6 across a wide range of Rad6 concentrations. The transfer of ubiquitin from Uba1 to Rad6 via transthioesterification (i.e., Rad6 ubiquitin thioester formation) was monitored under pre-steady-state (burst) conditions using 2.5 nM active Uba1 and the indicated concentrations of active Rad6, as described in the Experimental Procedures. (**A – E**) Quantitative analyses of Rad6 ubiquitin thioester conjugate formation. For each condition, the concentration of activated Uba1 (Ub~Uba1) and Rad6 ubiquitin thioester conjugates (Ub~Rad6) are plotted as a function of time. For the former (Ub~Uba1), all respective data points are fit to a flat line, yielding a y-intercept that is equal to the concentration of Ub~Uba1 (in nM) that is maintained throughout the respective experiment. For the latter (Ub~Rad6), all respective data points after time zero (i.e., t = 0) are fit to a linear regression yielding a y-intercept that provides an initial estimate of the amplitude of the observed burst (in nM) and a slope (m) that reflects the initial velocity ($v_{ss}$ in nM/s) of the steady-state phase. Data for each condition represent the average ± S.E.M. of at least three independent experiments. Panels **B** and **D** in the current figure are identical to **Figure 4A,** *Top* and *Bottom*, respectively, in the main text.
